# Adaptation-proof SARS-CoV-2 vaccine design

**DOI:** 10.1101/2022.05.17.492310

**Authors:** Yashavantha L. Vishweshwaraiah, Brianna Hnath, Brendan Rackley, Jian Wang, Abhinay Gontu, Morgan Chandler, Kirill A. Afonin, Suresh V. Kuchipudi, Neil Christensen, Neela H. Yennawar, Nikolay V. Dokholyan

## Abstract

Severe acute respiratory syndrome coronavirus 2 (SARS-CoV-2) surface spike glycoprotein - a major antibody target - is critical for virus entry via engagement of human angiotensin-converting enzyme 2 (ACE2) receptor. Despite successes with existing vaccines and therapies that primarily target the receptor binding domain (RBD) of the spike protein, the susceptibility of RBD to mutations provides escape routes for the SARS-CoV-2 from neutralizing antibodies. On the other hand, structural conservation in the spike protein can be targeted to reduce escape mutations and achieve broad protection. Here, we designed candidate stable immunogens that mimic surface features of selected conserved regions of spike protein through ‘epitope grafting,’ in which we present the target epitope topology on diverse heterologous scaffolds that can structurally accommodate the spike epitopes. Structural characterization of the epitope-scaffolds showed stark agreement with our computational models and target epitopes. The sera from mice immunized with engineered designs display epitope-scaffolds and spike binding activity. We also demonstrated the utility of the designed epitope-scaffolds in diagnostic applications. Taken all together, our study provides important methodology for targeting the conserved, non-RBD structural motifs of spike protein for SARS-CoV-2 epitope vaccine design and demonstrates the potential utility of ‘epitope grafting’ in rational vaccine design.

## Introduction

Coronavirus disease 2019 (COVID-19), caused by a novel coronavirus, severe acute respiratory syndrome coronavirus 2 (SARS-CoV-2), has caused a global health crisis^1^. Several safe and effective COVID-19 vaccines have been developed rapidly, emphasizing the importance of continued technological development efforts in vaccine development. However, viral evolution results in a continued need to expand the current vaccine repertoire by developing broadly protective, stable, safe, cost-effective, and scalable vaccine candidates for global use^2,3^.

SARS-CoV-2 belongs to the genus *Betacoronavirus* in the family *Coronaviridae*, which includes enveloped, single-stranded RNA viruses^4^. The surface spike protein of SARS-CoV-2 plays a crucial role in virus entry into cells by interacting with its receptor, the host angiotensin-converting enzyme 2 (ACE2) receptor^5^. Like other coronaviruses, the SARS-CoV-2 spike protein is a homotrimer composed of two functional domains, N-terminal S1 domain and C-terminal S2 domain^6^. The S1 domain mediates the ACE2 binding function through its receptor binding domain (RBD) and the S2 domain facilitates membrane fusion between the virus and the cell^7^. RBD has been widely targeted for the development of SARS-CoV-2 vaccines and monoclonal antibody-based therapeutics^8–10^. However, due to the error-prone nature of RNA-dependent RNA polymerase and the intense selective pressure exerted on the ACE2-interacting region of spike, RBD is more prone to mutations, leading to rapid adaptation and evolution of the virus^11–15^. Considering the critical role of RBD in binding SARS-CoV-2 to host cells, accumulation of mutations in this domain increases viral fitness by increasing ACE2 binding affinity or escaping neutralization, potentially resulting in loss of immunity in infected and vaccinated individuals. Accumulation of mutations in RBD alters the effectiveness of the RBD region-centered therapeutic antibodies and vaccines. Currently available SARS-CoV-2 vaccines have proven effective in reducing detectable symptomatic infections and complications of COVID-19. However, as transmission has progressed, several SARS-CoV-2 variants of concern^16^ with mutations in the RBD have emerged, leading to increased transmission and a reduction in neutralizing antibody response raised by viral infection or by currently approved vaccines. On the other hand, vaccines targeted at the conserved structural regions of the spike protein may overcome such limitations by combating adaptive capabilities of SARS-CoV-2, and could serve as the basis for universal vaccine development against variants of SARS-CoV-2 and various coronaviruses.

A number of different platforms and approaches are currently being used for vaccine development against COVID-19 including mRNA vaccines and adenoviral vector vaccines^17–19^. A rational, structure-guided vaccine design approach^20^ particularly aimed at eliciting the specific immune response to conserved viral epitope regions may provide broadly protective responses^21,22^. Epitope grafting or scaffolding is a high-precision approach, which has been effective in immunogen design by transferring conserved epitopes onto heterologous protein scaffolds for conformational stabilization^22–27^. Heterologous protein mimetics feature the epitope surface of the antigen and will be structurally compatible with the antibody binding mode.

Here, we sought to design immunogens for the SARS-CoV-2 spike protein using structure-guided epitope grafting and scaffolding methods. We recently described a general computational protocol to design epitope-scaffolds^20,27,28^. Using our established workflow^27^ (Fig. 1 and Supplementary Fig. 1), here we focused our protein design efforts on three distinct, conserved epitope regions in the S2 domain of spike protein to create structural mimics of the epitopes, and used these small protein mimics (epitope-scaffolds) as immunogens for a focused immune response. Biophysical and structural properties of the epitope-scaffolds indicated good agreement with our computational designs. Immunization with epitope-scaffolds induced a targeted immune response in mice against grafted epitopes. One of the epitope-scaffolds, whose solution structure was confirmed by small angle X-ray scattering, grafted with two different epitopes elicited a potent antibody response suggesting the inclusion of multiple epitopes in a single design for improved response. We also conceptually demonstrated the utility of epitope-scaffolds in serological applications for detecting epitope-specific antibodies. Overall, this study provides important methodology for utilizing the conserved structural elements of the spike protein in broadly protective immunogen design and the goal of developing a universal coronavirus (CoV) vaccine.

**Fig. 1.**
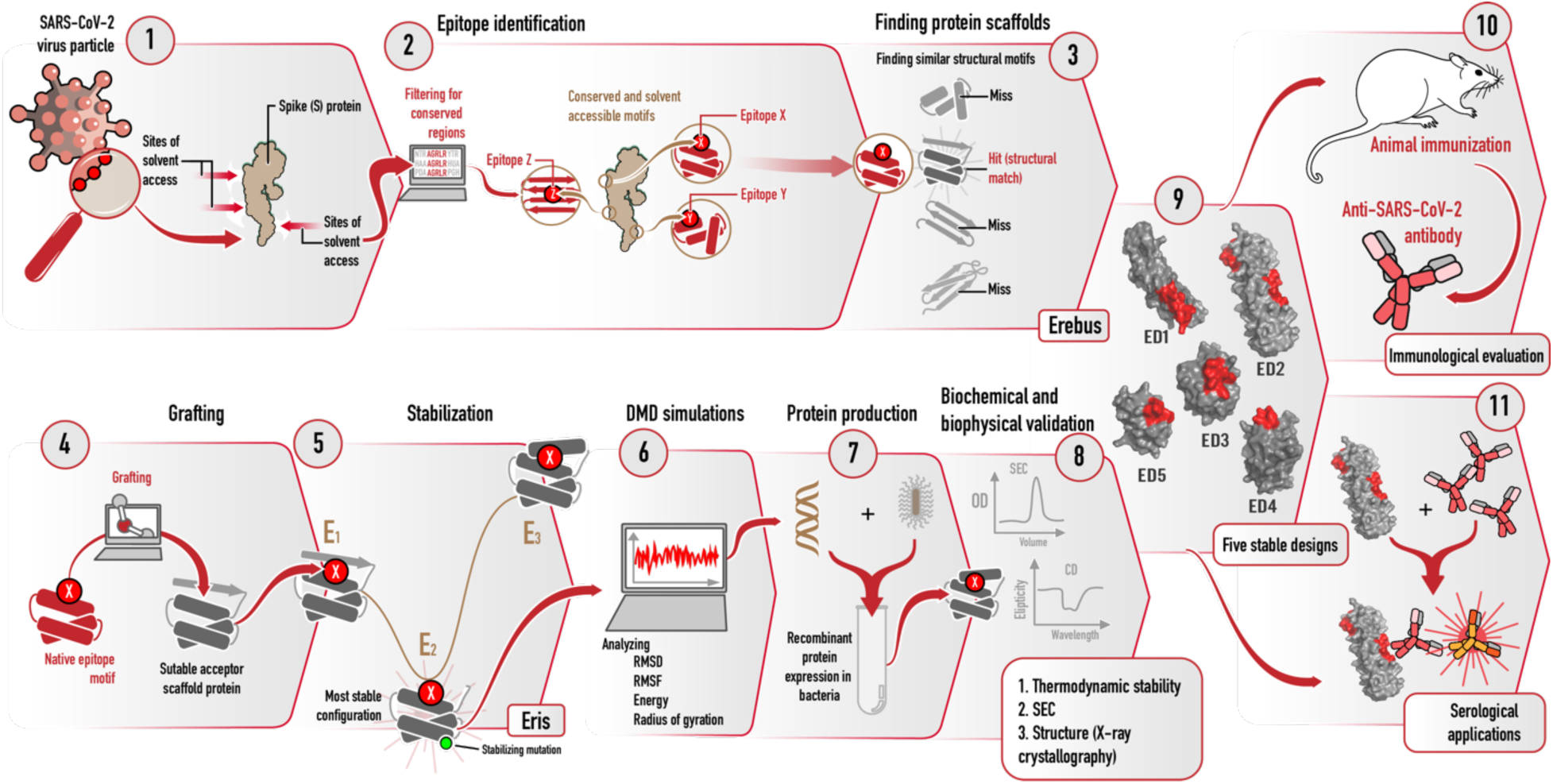
Schematic representation of different stages of the rational design of SARS-CoV-2 epitope-scaffold proteins for immune-focusing. Conserved and surface exposed spike epitopes were identified in steps 1 and 2. Suitable acceptor scaffold proteins were identified in step 3. Surface residues of the epitopes were grafted onto scaffold proteins (step 4) and stabilizing mutations incorporated (step 5). Confirmations of the grafted proteins were analyzed by DMD simulations (step 6). Stable proteins were expressed in *E*.*coli* and characterized by biophysical methods (steps 7 and 8). Five stable epitope-scaffolds (step 9) were tested for immunogenicity and serological applications (steps 10 and 11).

## Methods

### Computational design of epitope-scaffolds

We used ConSurf Web^29^ (http://consurf.tau.ac.il/) to calculate the evolutionary conservation score for each amino acid of spike protein (PDB: 6VSB). The conservation of epitopes was analyzed by multiple sequence alignment with Clustal Omega^30^. We analyzed the solvent accessible area on protein surface by using Gaia^31^ and Chiron^32^. We used the standalone version of Erebus^33^, a substructure search server (https://dokhlab.med.psu.edu/erebus), to identify scaffold proteins for grafting. We define the query structure for Erebus as the backbone atoms (Cα, N, C=O) of each spike-epitope. We extracted the coordinates of backbone atoms from the PDB structure of 6VSB for the identified three spike-epitopes. We provided the query structures (PDB format) to Erebus with default search parameters (matching precision 0.3 Å). Erebus exhaustively searched the PDB database for matches of structural scaffolds to atom pairs in the query. Erebus results in output scaffolds ranked based on their root-mean-square deviations (RMSD) to the query structure. We redesigned the scaffolds using Eris^34,35^ (https://eris.dokhlab.org), a computational platform that automatically performs side-chain repacking and backbone relaxation and calculates the changes in free energy upon mutations (ΔΔG = ΔG_mut_ – ΔG_wt_)^34,35^. To support massive computational mutagenesis, we used the standalone version of Eris. We introduced the solvent-exposed residues within each epitope into the corresponding scaffolds by Eris. For every single mutation, we performed 1000 Eris calculations to reach a converged distribution of ΔΔG values. We obtained the average ΔΔG and its standard deviation. We discarded destabilizing mutations (ΔΔG > 0 kcal mol^-1^). We optimized the chimeric structures using Chiron and Gaia. To estimate the structural rigidity of design models, we performed discrete molecular dynamics (DMD) simulation^36–39^, an event-driven simulation that employs a discrete potential energy that relies on the calculation of atomic collisions. We performed three independent simulations for each design, and each simulation was performed for 10^7^ steps. Each step in DMD corresponds to approximately 50 ps. We analyzed the trajectories by using MDAnalysis to derive information about the average structure, RMSD, energy, radius of gyration, and root-mean-square fluctuations (RMSF).

### Protein expression and purification

We obtained the genes encoding the designed proteins in pET28a vectors from GenScript (Genscript.com). We confirmed the sequences of all the constructs by DNA sequencing (Eton bioscience, USA). We transformed each individual construct into *E. coli* BL21 (DE3) pLysE strains and cultured them to an optical density of 0.6-0.8 at 600 nm in LB broth. We induced the gene expression with 1 mM isopropyl-β-d-thiogalactopyransoide (IPTG). We purified the N-terminal His-tagged proteins on a HisTrap column (GE Healthcare) and then by size exclusion chromatography (SEC) (Superdex S-200preparatory-grade column (GE Healthcare)). Briefly, we resuspended cell pellets in lysis buffer (20 mM Na_2_HPO_4_, 40 mM imidazol, 500 mM NaCl, pH 7.4) with protease inhibitors (1 mM phenylmethylsulfonyl fluoride and 1 µM pepstatin A) then lysed by sonication. We separated the supernatant containing protein components from precipitate by centrifugation at 15000 rpm for 30 min at 4°C. After passing through 0.22-μm filter (Millipore), we loaded the supernatants onto a 5 mL HisTrap HP column (GE Healthcare) using an AKTA Pure FPLC machine. We eluted the proteins with 500 mM imidazol in 20 mM Na_2_HPO_4_, 500 mM NaCl, and pH 7.4. We further purified the proteins through a 25 mL Superdex 200 preparatory-grade column (GE Healthcare) with 20 mM Na_2_HPO_4_ and 100 mM NaCl (pH 7.4) as the running buffer. We determined the protein concentrations using BCA assay kit (ThermoFisher Scientific).

### Size exclusion chromatography

We loaded the proteins (1 mg/mL protein in 20 mM Na_2_HPO_4_, 100 mM NaCl, pH 7.4) onto a Superdex 200 column (GE Healthcare) at a flow rate of 0.5 mL/min using an AKTA Pure FPLC machine. We monitored UV absorption at 280 nm. We ran standards (Gel filtration standard, Bio-Rad) on the cleaned column prior to running the samples. The standards included five molecular weight markers: vitamin B12 (1.35 kDa), myoglobin (17 kDa), ovalbumin (44 kDa), γ-globulin (158 kDa), and thyroglobulin (670 kDa). We calculated the apparent molecular weight of designed proteins by plotting log of molecular weight versus elution volume for all the proteins.

### Circular Dichroism (CD) Spectroscopy

CD measurements were performed on a JASCO J-1500 spectrometer, equipped with a Peltier model PTC-517 thermostat cell holder. We used a sample concentration of 0.2 mg/ml. Signals were recorded from 260 nm to 185 nm with a scan speed of 50 nm/min and a band width of 1 nm at 20 °C. The quartz cell used was 1 mm in length. Samples were scanned three times and the scans were averaged. For all spectra, a buffer reading was subtracted. The secondary structures of proteins were compared using the Jasco spectra manager software. For melting curves, temperature-induced protein denaturation was followed by a change in ellipticity at 210 nm.

### X-ray Crystallography

The crystals of ED1 epitope-scaffold were grown using the vapor diffusion method with hanging drops. We mixed a solution consisting of 0.2 M sodium formate and 20% (w/v) PEG 3350 crystallization solution 1:1 with a protein solution concentration of 10 mg/ml in 20 mM Tris-HCl at pH7.4. We set up the crystallization trials at 10 °C. Needle crystals appeared in 21 days and grew to full size in 4 weeks. For cryo freezing crystals, we soaked them in mother liquor with 25% glycerol for 60 min prior to X-ray data collection. We collected the X-ray data on the Rigaku Micromax 007HF rotating anode microfocus X-ray source equipped with Oxford cryo-stream, universal 4-circle kappa goniometer, Varimax VHF optics and a HyPixArc150 detector. X-ray Diffraction data set to a resolution of 1.99 Angstroms were collected and processed using the Rigaku CrysAlis Pro software. We solved the crystal structure by the molecular replacement method using the apolipoprotein E amino-terminal domain structure (1BZ4) as a search model in the PHENIX software. Several iterations of model building in COOT program and refinement in the PHENIX program were done to complete the structure. Pymol software was used for all the structural analysis and figures.

### Small angle X-ray scattering and modeling

We collected the BioSAXS data for the ED2 epitope-scaffold protein on the in house Rigaku BioSAXS2000^nano^ equipped with a Kratky camera system and housed on a MicroMax007 X-ray generator. The system includes OptiSAXS confocal max-flux optics that is designed specifically for SAXS/WAXS data and a sensitive HyPix-3000 Hybrid Photon Counting detector. The protein samples were ultracentrifuged in 20 mM Tris-HCl at pH7.4 and 150 mM NaCl at 100,000 rpm for 20 min for 2, 4, and 8 mg/mL concentrations and these SAXS data sets showed a concentration dependent multimerization. A subsequent SAXS run with freshly eluted protein sample from an SEC run at a concentration of 0.65 mg/ml in 20 mM Tris-HCl, pH 7.4 and 500 mM NaCl was used, and data was collected in triplicates. The sample was picked by an autoloader into a quartz flow-cell mounted on a stage cooled to 4 °C and aligned in the X-ray beam. We collected six ten-minute scattering images, checked for any radiation damage and averaged. The triplicate sample data sets overlaid well and were further averaged. We also collected the buffer data sets similarly for reference subtraction to get the SAXS scattering curve from only the protein. Data collection and processing was done using the Rigaku SAXSLab software. We subsequently analyzed the data files for radius of gyration, Guinier fit, Kratky plot, and pair distance distribution function using the ATSAS software. We subsequently used the program DAMMIN, part of ATSAS suite, for *ab initio* low resolution shape determination. Solvent envelopes were also computed using DENSS, an algorithm used for calculating *ab initio* electron density maps directly from solution scattering data. A dimer model was generated by the SASREF refinement program by inputting two subunits of the four-helix bundle (4UOS) and the experimental SAXS curve. Subsequently, energy minimization of the dimer was done using the YASARA server. The CRYSOL program was finally used to compare the experimental and calculated SAXS profiles.

### Immunization of mice and rabbits

All animal studies were reviewed and approved by the Penn State University College of Medicine IACUC (PROTO201900719). Animal studies followed guidelines from the NIH regarding the care and use of animals in research. Mice (strain FVB) were immunized by standard methods using purified epitope-scaffolds admixed with sigma adjuvant (RIBI adjuvant) at a 1:1 ratio as described by the manufacturer. Animals were immunized with 100-200 µg protein in a volume of 100 µL of adjuvanted preparation by i.p. immunization of mice and 200 µL for i.m. immunization of rabbits. Booster immunizations (2 per animal) were conducted 2-3 weeks after each immunization with a final booster in saline. Animals were anesthetized prior to the collection of blood samples and euthanized for terminal harvesting of tissues and blood.

Monoclonal antibodies were prepared by standard hybridoma fusion technology with polyethylene glycol as previously described^40^. The fusion partner for mouse hybridomas was P3XAg.8.683 (ATCC). Fusion reactions were plated into 96-well flat-bottomed cell culture plates and hybridomas selected using standard hypoxanthine/aminopterin/thymidine selection methods. Growing hybridomas were marked on the wells of the plates and supernatants of these wells tested in an enzyme-linked immunosorbent assay (ELISA) for reactivity to the ED2 construct as described below. Positive wells were selected for cloning and retesting in ELISA and neutralization assays.

### Enzyme-linked immunosorbent assay (ELISA)

We tested the binding properties of antibodies to the epitope-scaffolds and spike protein using in-house ELISA. We coated ninety-six-well ELISA plates (Nunc MaxiSorp, Thermo Fisher Scientific) overnight with epitope-scaffolds in PBS buffer. We included the native scaffolds (non-transplanted) as controls to confirm the specificity of epitope-scaffolds. We blocked the plates with 100 μL of 1% BSA in PBS buffer and we incubated for 2 h at room temperature. We tested the binding of immunized mouse serum samples, COVID-19 patient’s serum samples (1:200 diluted) and commercially obtained anti-spike antibodies against the coated proteins. After incubating at room temperature for 2 h, we extensively washed the plate with PBS containing 0.1% Tween-20. We added horseradish peroxidase (HRP)-conjugated goat anti-human IgG (1:5000, SouthernBiotech #2016-05) to detect the binding in COVID-19 patients’ serum samples, HRP-conjugated goat anti-mouse IgG (1:5000, Sigma-Aldrich, #AP124P) to detect the binding in mouse serum samples, HRP-conjugated goat anti-rabbit IgG (1:5000, ThermoFisher Scientific, #31460) to detect the binding in commercial anti-spike antibodies, for 1 h at 37 °C. We then washed the ELISA plates with PBS containing 0.1% Tween-20. Subsequently, we added 100 μL of HRP substrate (3,3′,5,5′-Tetramethylbenzidine (TMB), Sigma-Aldrich) into each well. Immediately we stopped the reaction by adding 50 μL of 2 N H_2_SO_4_ solution and analyzed on a microplate reader (Molecular Devices, Spectramax I3) at 450 nm wavelength. We obtained the normalized results by calculating the difference between the OD of the protein-coated well and the BSA-coated well. For the COVID-19 samples, we obtained the graft-specific response by calculating the difference between the OD of the ED2-coated well and the ED2 control-coated well. Culture supernatants from wells containing hybridomas were initially screened for reactivity to epitope-scaffolds and native scaffolds attached to ELISA plate wells using alkaline buffer^40^. Positive cultures were expanded, cloned and the clones retested for reactivity, again expanded, and stored as aliquots for various assays (ELISA and neutralization). Dilutions of culture supernatant were titrated in the ELISA to establish half maximum binding activity (dilution of supernatant that produced 50% of maximum O.D. values).

### ELISA based on ED2-decorated magnetic nanoparticle (MNPs)

To conjugate ED2 epitope-scaffold proteins to magnetic beads, a DNA duplex with biotin and nitrilotriacetate (NTA) on opposite 5’ ends was used as a linker between streptavidin-decorated MNPs and the His6-tags present on ED2. Streptavidin-decorated MNPs (RayBiotech, Inc. #801-106-1) were transferred (50 µL of 5 mg/mL) to a tube, pulled down for two minutes on a magnet, and the supernatant was replaced with 500 µL of 1.27 µM biotinylated DNA (5’-/5Biosg/CGGTGGTGCAGATGAACTTCAGGGTCA from Integrated DNA Technologies, Inc.) for a 1 MNP : 100,000 DNA ratio in 25 mM Tris-HCl, 137 mM NaCl, 2.68 mM KCl, pH 7.4 (1X tris-buffered saline, TBS) as previously described^41^. After one hour of incubation at RT, the MNPs were again pulled down for two minutes on a magnet, resuspended in 500 µL of 1X TBS with 0.05% Tween-20 (1X TBST) which was then removed as a washing step, and then brought up in 500 µL of 1.27 µM NTA DNA (5’-[NTA][SS-C6]ACCCTGAAGTTCATCTGCACCACCG from GeneLink) in 1X TBS. Following one hour of incubation at RT, the MNPs were once more pulled down, washed with 1X TBST, and the supernatant was replaced with 500 µL of 1X TBS with 2.54 µM NiCl_2_, followed by one hour of incubation at RT. The MNPs were pulled down and resuspended in 1X TBS prior to all experiments. We collected the ED2-decorated MNPs (later referred to as ED2-MNPs) by subjecting the mixture to a magnetic field and washing three times with PBS. We estimated the protein concentration using BCA assay kit (ThermoFisher Scientific). To demonstrate the binding, we first mixed the diluted serum (1:100, 1:1000, and 1:10000) samples (in PBS) with 250 ng of proteins (ED2-MNPs) in a microtiter plate and incubated for 30 min at room temperature. We washed with PBS three times to remove the unbound fractions and treated the mixtures with anti-human IgG secondary antibody (SouthernBiotech #2016-05). The plate was further incubated at room temperature for 30 min, and then washed with PBS three times by placing the ELISA plate over an in-house magnetic plate. We then added 100 μL of TMB into each well. We immediately stopped the reaction by adding 50 μL of 2N H_2_SO_4_ solution and measured the absorbance on a microplate reader (Molecular Devices, Spectramax I3) at 450 nm wavelength.

### Live Virus Neutralization (VN) assays

We evaluated the ability of epitope scaffolds and spike protein antibodies to neutralize SARS-CoV-2 using live virus neutralization (VN) assays that we previously described^42–45^. Briefly, serum samples were diluted twofold in triplicate and incubated with 100 tissue culture infective dose 50 (TCID50) units of SARS-CoV-2 strain USA-WA1/2020 (NR-52281-BEI Resources, USA) at 5% CO_2_ at 36 ± 2 °C for 60 min. The antibody–virus mixture was then added to monolayers of Vero E6 cells (CRL-1586, ATCC, USA) in 96-well microtiter plates and incubated further for 72 h at 5% CO_2_ at 36 ± 2 °C. After staining the plates with crystal violet formaldehyde stain for one hour, the plates were visually inspected for cytopathic effect (CPE) or protection. The reciprocal of the highest dilution of the plasma where at least two of the three wells were protected (no CPE) was determined as the VN titer of the sample. All the experiments with live SARS-CoV-2 were performed under BSL-3 containment condition in the Eva J Pell laboratory for advanced biological research at Penn State.

### Ethical approval and patient’s sera

We obtained the COVID-19 positive human samples and healthy human serum samples (pre-pandemic samples) from Penn State College of Medicine Institute for Personalized Medicine repository. All the human samples were obtained under protocols approved by the Penn State Health Institution Review Board.

### Statistics and reproducibility

We performed all statistical analyses by unpaired two-tailed Student’s t-test using GraphPad Prism 8 v8.2.1 software and Microsoft Excel v16.49. The levels of significance are denoted as *P ≤ 0.05, **P ≤ 0.01, ***P ≤ 0.001, ****P ≤ 0.0001, and NS not significant (P > 0.05). We used GraphPad Prism 8 version 8.2.1 and Adobe Illustrator to draw and assemble the figures.

## Results

### Computational design of epitope-scaffolds

We identified three epitope regions on the S2 domain of spike protein of SARS-CoV-2 based on the previously determined trimeric spike structure^46^ (Fig. 2a and Supplementary Fig. 2). The rationales for identifying these epitopes were: (i) high sequence conservation and (ii) high solvent-exposed area on the spike surface. All the residues in epitope-1 and epitope-2 are located on α-helices (epitope-1, residues 921-936 and epitope-2, residues 816-824) and the residues in epitope-3 are located on a loop region (residues 789-797). To design immunogens that mimic the three epitopes, we first searched the entire Protein Data Bank (PDB) for suitable acceptor proteins (scaffolds) that share backbone structural similarity with identified epitopes. We used a rapid structural motif-mining algorithm *Erebus* to identify suitable potential scaffolds. We estimated the surface matching between scaffolds and epitopes by analysis of root-mean-square deviations (RMSD, 0.5-3.0 Å) and selected the top 30 ‘hits’ for each epitope. Using a side-chain grafting approach^28^, we replaced the surface amino acids of the epitope match region within each scaffold with corresponding residues in the consensus sequences of spike epitopes. For each epitope, we created at least 15 epitope-scaffold designs and calculated the change in free energy of the substitutions (ΔΔG_mut_) using *Eris*. We applied a filter by excluding the designs having highly destabilizing mutations (ΔΔG_mut_ > 4 kcal mol^-1^). We further optimized the epitope-scaffolds by introducing stabilizing mutations into each scaffold (ΔΔG_mut_ < 0 kcal mol^-1^). We evaluated the conformational stabilities of all the epitope-scaffolds using DMD simulations and selected the epitope-scaffolds that have substantial rigidity around the grafted region (Supplementary Fig. 3). Overall, we selected seven epitope-scaffold designs for three epitopes based on RMSF and RMSD analysis, radius of gyration, and energy analysis (Supplementary Fig. 3), which we named ED1-ED7 (Supplementary Table 1). Except for ED2, all other epitope-scaffold designs were grafted with a single epitope. ED2 was grafted with epitope-1 and -3 at different sites. For comparison, we also included corresponding native scaffolds (non-transplanted) in the simulations.

**Fig. 2.**
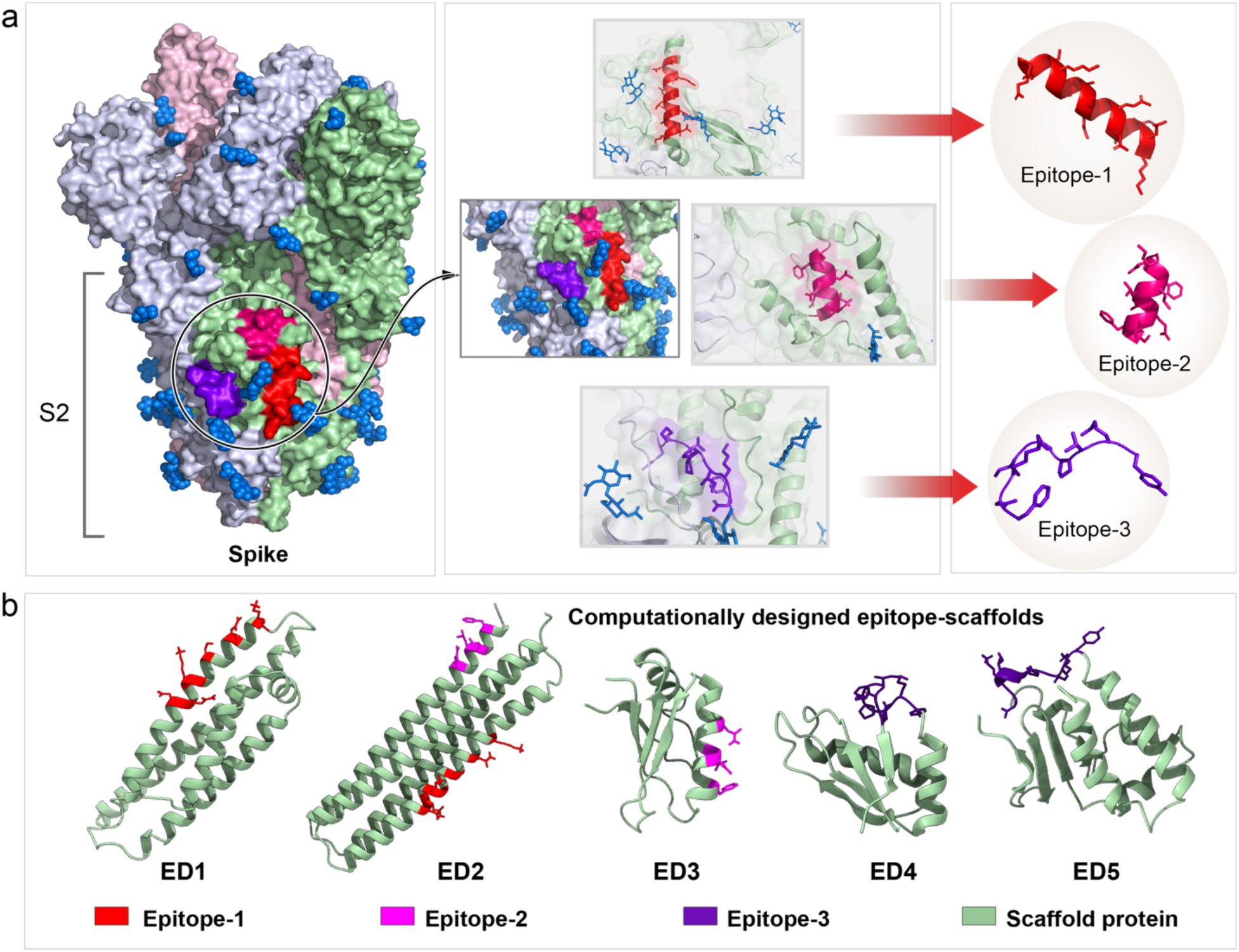
Identification of SARS-CoV-2 epitopes and computational designs of epitope-scaffold proteins. **a**, Three epitopes from the S2 domain of spike were selected based on sequence conservation and surface exposure. Three epitopes are shown in red, pink, and purple color. **b**, Computational models of epitope-scaffolds. Five stable, computationally designed epitope-scaffolds were expressed in *E*.*coli*. Scaffold proteins are shown in green. Grafted regions are shown in red, pink and purple.

### Biophysical and structural characterization

We expressed seven chimeric immunogens (epitope-scaffolds) and their corresponding native scaffolds in *E. coli*, purified by affinity chromatography and size exclusion chromatography. We purified five epitope-scaffolds (ED1-ED5, Figure 2b) and all the native scaffolds as soluble proteins; we observed aggregation for one protein (ED4) (Supplementary Table 2). SEC data suggested a monomeric conformation for all five epitope-scaffolds (Fig. 3a). The five epitope-scaffolds displayed CD spectra consistent with the designed topology (Fig. 3b). Secondary structure comparison between the epitope-scaffold designs with native scaffolds showed minimal changes. We measured the stabilities of all the designed proteins’ thermal denaturation using CD spectroscopy melting curve analysis. The melting curve analysis showed a melting temperature (T_m_) in excess of 90 °C for ED2, ED3, and ED4, whereas for ED1 and ED5, T_m_ was in the range of 46-52 °C (Supplementary Table 3).

**Fig. 3.**
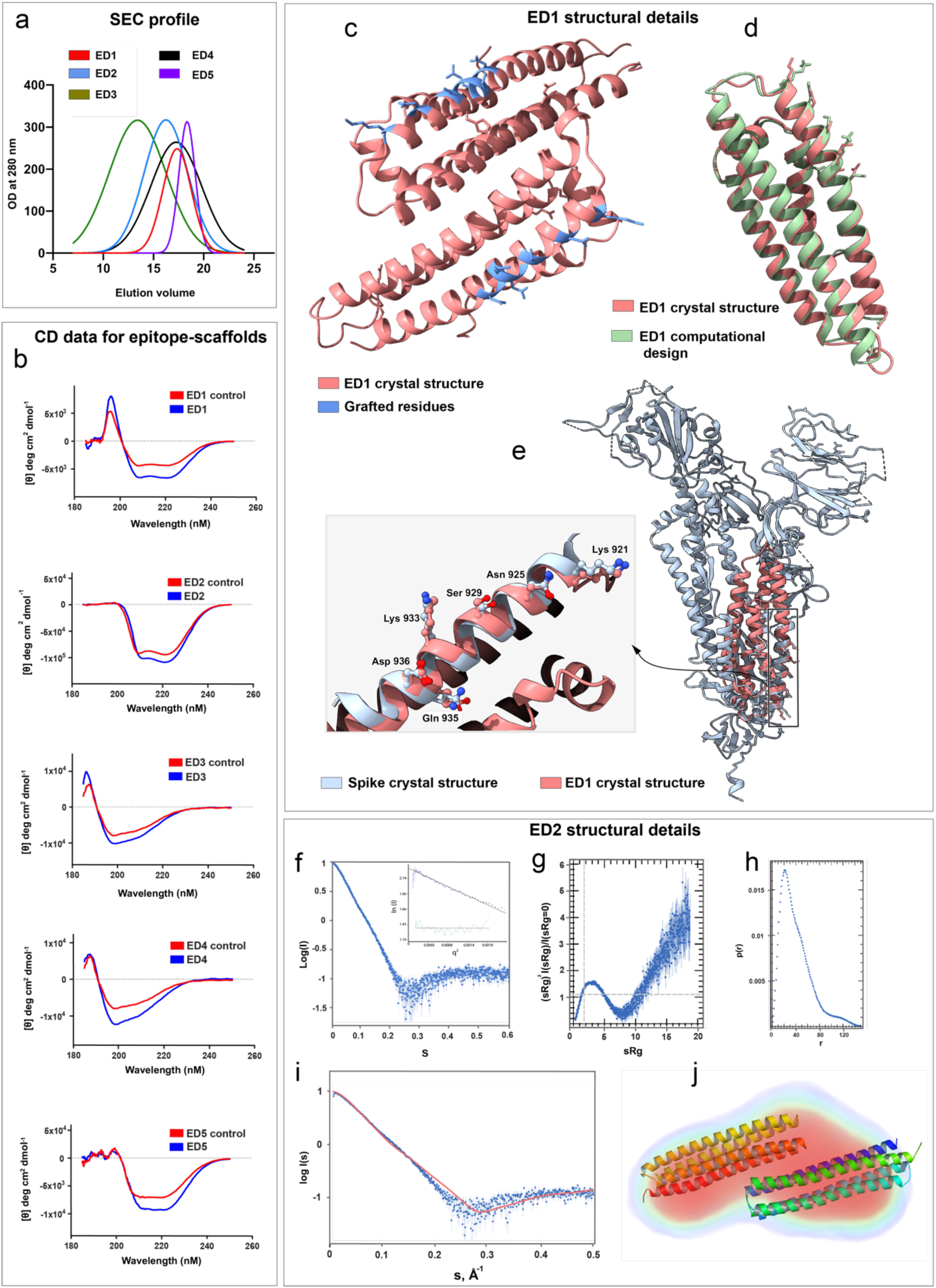
Structural characterization of designed epitope-scaffolds. **a**, Analytical SEC profile for the five epitope-scaffolds. Of the five designs, four folded into monomers and ED2 formed a dimer in solution. **b**, The CD spectra of five epitope-scaffolds compared to native scaffolds (controls). CD data indicated that the expected secondary structures were retained in solution for all designs. **c**, Crystal structure of ED1, with the grafted residues shown in blue. **d**, Crystal structure of ED1 aligned to the computational design model. ED1 is shown in salmon color and the design model is shown in green. Epitopes are shown as sticks. **e**, Crystal structure of ED1 aligned to the corresponding epitope region on spike protein. The enlarged image shows the close-up view of the alignment between grafted residues in ED1 (salmon) and the corresponding spike epitope residues (blue). Epitopes are shown as ball and sticks. **f**, SAXS raw data for ED2 at 0.7 mg/ml in 20 mM Tris-HCl at pH7.4 and 500mM NaCl. Inset shows the Guinier Plot for the ED2 data. **g**, Kratky plots derived from the SAXS data to qualitatively assess the flexibility and/or degree of unfolding in protein. The results indicate a well folded protein. **h**, Pair distance distribution function (p(r)) analysis of ED2. P(r) analysis suggests dimeric ED2 in solution. **i**, The CRYSOL program was used to the calculate SAXS profile of the dimer (red) on the experimental SAXS profile (in blue) (Chi-square fit of 6.8). **j**, The DENsity from Solution Scattering (DENSS) electron density map shown as a transparent surface fits a dimer of ED2. The dimer was generated by the SASREF refinement program and energy minimized using the YASARA server.

To evaluate the epitope structural mimicry, we solved the crystal structure of ED1 at 2.0 Å resolution (Table 1). The structure of ED1 revealed a dimer in contrast to the monomer observed in SEC (Fig. 3c). Experimental structure of ED1 featured remarkable similarity to the ED1 computational model; ignoring the two short loop regions that connect the helices, Cα RMSD between the computational model and the experimental structure was 0.8 Å (Fig. 3d and Supplementary Fig. 4). Comparison with spike protein revealed that all the grafted residues in the ED1 experimental structure largely recapitulated the conformations of the corresponding residues on spike protein. Within the epitope regions, Cα RMSD between the ED1 experimental structure and spike protein was 0.6 Å, suggesting a high degree of mimicry (Fig. 3e).

**Table 1.** Crystallographic data collection and model refinement statistics for ED1.

|  |  |
| --- | --- |
| *Data Collection* |  |
| Wavelength (Å) | 1.54184 |
| Frames/Runs | 1148/4 |
| Rotation (°) | 360 |
| Space group | P 2<sub>1</sub> |
| *Unit cell dimensions* |  |
| a, b, c (Å) | 39.060(2), 95.631(3), 40.916(3), |
| α, β, γ (°) | 90.0, 108.576(8), 90.0 |
| Resolution (Å) | 95.63 - 1.99 |
|  | (2.06 - 1.99)<sup>a</sup> |
| R<sub>int</sub><sup>b</sup> | 0.111 (0.216) |
| I/σ(I) | 14.6 (2.6) |
| Completeness (%) | 97.0 (94.3) |
| Average redundancy | 3.8 (3.5) |
| *Structure refinement* |  |
| Resolution range (Å) | 17.87-1.99 |
| No. unique reflections | 18932 |
| R<sub>work</sub><sup>c</sup>/R<sub>free</sub><sup>d</sup> | 0.262/0.292 |
| *Number of atoms* |  |
| Protein | 2670 |
| Ligand/ions | 4 |
| Water | 277 |
| *Avg. B-factor* |  |
| Protein | 20.04 |
| Ligand/ions | 22.7 |
| Water | 20.3 |
| *RMS deviations<sup>e</sup>* |  |
| Bond lengths (Å) | 0.002 |
| Bond angles (°) | 0.44 |
| *Ramachandran plot<sup>f</sup>* |  |
| favored (%) | 93.8 |
| allowed (%) | 4.5 |
| outliers (%) | 2.4 |

<sup>a</sup>The highest resolution shell is shown in parenthesis
<sup>b</sup> $R_{int} = \sum_{hkl} \sum_i |I_i(hkl) - \langle I(hkl) \rangle| / \sum_{hkl} \sum_i I_i(hkl)$ , where $I_i(hkl)$ is the intensity of an observation and is the mean value for its unique reflection. Summations are over all reflections.
<sup>c</sup> $R_{work} = \sum_h |F_o(h) - F_c(h)| / \sum_h F_o(h)$ , where $F_o$ and $F_c$ are the observed and calculated structure factor amplitudes, respectively.
<sup>d</sup> $R_{free}$ was calculated with 5% of the data excluded from the refinement.
<sup>e</sup>RMS deviation, root mean square deviation from ideal values.
<sup>f</sup>The categories were defined by Molprobit in PHENIX.

We also obtained structural information for ED2 design using BioSAXS (Fig. 3f-j and Supplementary Table 4). For ED2, we observed multimerization at higher concentration (concentration > 5-7 mg/mL). In contrast to our SEC data, SAXS data suggested a dimeric conformation of ED2 in solution. The Guinier fit suggested a radius of gyration of 30.6 (+/-0.2) Å. The Kratky plot to assess the flexibility and/or degree of unfolding suggested a well-folded protein. Pair distance distribution function *P*(r) peaked at 33.5 Å, in agreement with the Guinier analysis, and suggested a maximum distance of 150 Å. The *P*(r) analysis suggested a dimer in solution as the maximum distance for a monomer is around 80 Å.

### Immunogenic evaluation

Our analysis of immunogenicity for four stable epitope-scaffolds (ED1, ED2, ED3, and ED5) in mouse models revealed epitope-specific antibodies. ED1 and ED3 designs induced low levels of epitope-specific IgG response as determined by enzyme-linked immunosorbent assay (ELISA) (Fig. 4a). In contrast, ED2 immunized mice groups showed markedly higher level of antigen specific IgG response, with the Optical Density (OD) measurement of ∼1. ED5 immunized animals showed a moderate response with the OD measurement of ∼0.5. Based on the antibody response, we selected ED2 antisera for further evaluation. ELISA comparison of binding of ED2 antisera against ED2 epitope-scaffold and its native control scaffold showed a graft-specific IgG response (Fig. 4b). When we further tested the ED2 antisera against spike protein, antisera displayed significant binding (Fig. 4b). These results prompted us to generate a panel of monoclonal antibodies (mAbs). We selected the mice which had the highest titer to ED2 epitope-scaffold for fusion. We selected the B-cell hybridoma clones based on ED2 epitope-specific binding. We generated a panel of 40 mAbs in this manner, selected a total of 21 ED2 epitope-scaffold specific mAbs, and tested them against spike protein (Supplementary Fig. 5). MAbs failed to show any response against spike protein (Supplementary Fig. 5). We then tested the ED2 epitope-scaffold induced mouse antisera and the screened MAbs for live SARS-CoV-2 virus neutralizing activities. Mouse antisera showed a limited neutralizing effect (neutralization activity in one out of three wells, data not shown). None of the MAbs displayed live SARS-CoV-2 virus neutralization activity.

**Fig. 4.**
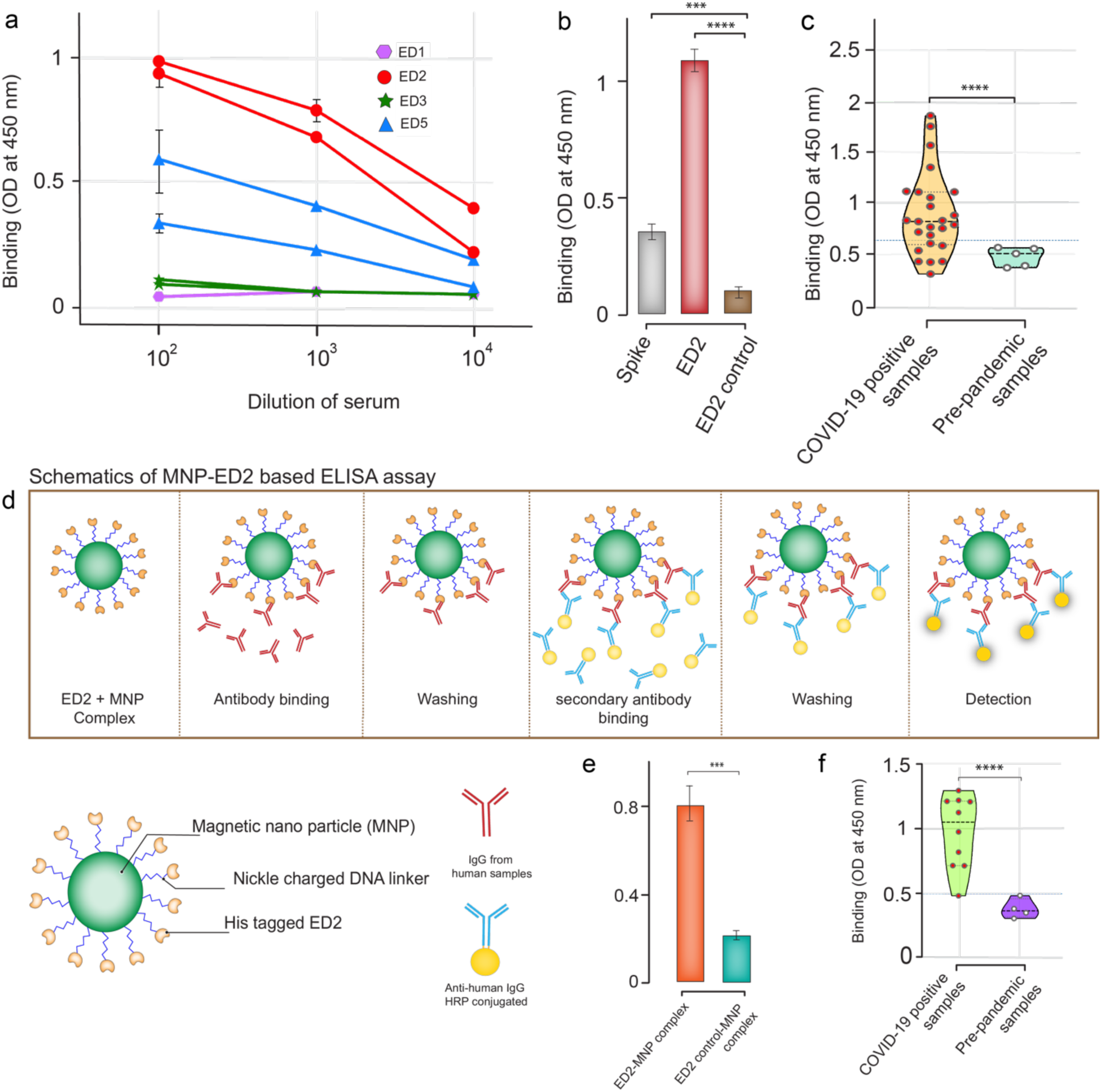
Immunological evaluation and serological application of epitope-scaffolds. **a**, Sera from mice immunized with epitope-scaffolds were tested by ELISA for binding to each epitope-scaffold. Data include two animals per group. **b**, Comparison of ED2 immunized mice serum response to ED2, ED2 control, and spike protein. **c**, Anti-epitope-1 IgG antibodies were detected in both negative controls (pre-pandemic samples) and COVID-19 patient samples using ED2. Data represent violin plots and individual data points. n = 5 samples for pre-pandemic samples and n = 25 samples for COVID-19 patient samples. **d**, Schematic representation of ELISA based on ED2-decorated magnetic nanoparticle (MNPs) for the detection of epitope-1 specific IgG antibodies. **e**, Anti-epitope-1 IgG antibodies were tested in negative controls and COVID-19 patient samples using ED2 and by employing MNP-based ELISA. **f**, Anti-epitope-1 IgG antibodies were tested in negative controls and COVID-19 patient samples using ED2 and by employing MNP-based ELISA. Data represent violin plots and individual data points. n = 4 samples for pre-pandemic samples and n = 10 samples for COVID-19 patient samples.

We evaluated the ED epitope-scaffolds for binding to commercially available anti-spike polyclonal antibody using in-house indirect ELISA assay by coating the epitope-scaffolds. We included the native scaffold proteins to measure the background binding. Of the five epitope-scaffolds tested, ED2 and ED5 demonstrated significant binding, with ED2 displaying a high OD readout (Supplementary Fig. 6).

To obtain insights on whether humans can produce antibodies specific for ED2 scaffolds, we tested the binding of sera from 25 SARS-CoV-2 positive humans to ED2 epitope-scaffolds using our in-house IgG ELISA assay (all samples were of unknown time from symptom onset). We included 5 healthy human samples as controls (samples collected prior to the SARS-CoV-2 pandemic). We observed significant binding in SARS-CoV-2 positive samples compared to the healthy samples (Fig. 4c). Results also indicated a clinically relevant conformation of the epitope-scaffold, suggesting potential application of epitope-scaffolds in diagnostics, especially in identifying epitope-specific antibodies.

We further used ED2 epitope-scaffold proteins to conceptually demonstrate a rapid ELISA based on ED2-decorated magnetic nanoparticles to assess the epitope-structure-specific antibodies in polyclonal sera (Fig. 4d). We conjugated the histidine-tagged ED2 epitope-scaffold proteins to nickel charged magnetic nanoparticles and first analyzed binding against commercial anti-spike polyclonal antibodies (Fig. 4e), then we tested against ten SARS-CoV-2 positive samples and 4 healthy human samples (Fig. 4f). The ten SARS-CoV-2 positive samples tested here were selected from the ELISA results of our in-house assay. Using this approach, we were able to complete the ELISA results for the tested samples in less than 60 min. For the samples tested, our magnetic nanoparticle-based liquid-ELISA showed a similar trend in absorbance compared to the in-house ELISA (Supplementary Fig. 7).

## Discussion

The majority of SARS-CoV-2 vaccines currently being developed or deployed target the spike RBD as their main antigenic component^47^. Although RBD is a more attractive target, RBD is a less conserved region of spike protein and more prone to accommodate mutation. Multiple mutations within the RBD are a major cause of the constantly emerging SARS-CoV-2 variants and VOCs. These mutations are reported to significantly reduce the effectiveness of some currently approved vaccines such as Pfizer/BioNTech BNT162b2, Moderna mRNA-1273, or ChAdOx1^48–53^. Also, it is still unclear about the breadth of protection provided by the currently approved vaccines against sarbecoviruses and other CoVs. These data point to the urgent need to develop pan-sarbeco vaccines or universal CoV vaccines to protect against emerging SARS-CoV-2 variants and future CoV outbreaks.

The primary requirement for a universal vaccine is the achievement of broad immunity^54–56^. Eliciting a focused immune response against the conserved epitopes of the spike could induce broad immunity^22,27^. Broadly effective SARS-CoV-2 immunogens targeting the conserved regions of spike are expected to effectively block current and emerging neutralization-resistant and highly infectious SARS-CoV-2 variants that evade the antibody neutralization response by mutating key residues on the spike protein. Such immunogens are expected to be effective against other CoVs as well. Conventional vaccine approaches have not been successful in producing broadly effective vaccines for highly variable viral pathogens^57,58^. While the structure-guided antigen design approach not only offers speed and precision for vaccine design^20,59^, in combination with the epitope grafting or scaffolding method, this approach has shown a great promise in epitope vaccine design against highly variable viral pathogens such as HIV-1 and RSV^23–25,27,58^. In the present study, we reported the design of candidate immunogens by targeting the conserved structural regions of spike protein for a focused immune response against SARS-CoV-2 through structure-guided engineering and grafting approaches.

Our approach utilizes the atomic-level details from the spike crystal structure to design immunogens that mimic the conformation of largely neutralizing, non-RBD conserved epitopes on the spike protein surface. The elicitation of structure-specific antibodies against SARS-CoV-2 using grafting has recently been reported by targeting the moderately conserved region of RBD^60^. As RBD is more susceptible to intensive mutations, structural conservation in non-RBD regions, such as the S2 domain, could be exploited for vaccine design to achieve broader immunity; S2 domain is more conserved across CoVs. Cross-reactive antibodies, nAbs, and CD4+ T cells are reported to target the S2 domain of SARS-CoV-2 and other CoVs^61–67^, suggesting that S2 is a vaccine target. We grafted three conserved epitope regions from the S2 domain of SARS-CoV-2 onto antigenically distinct scaffold proteins to produce small, stable, and soluble proteins that closely mimic the target epitopes. Biophysical and structural characterization data validated our computational designs. The immunization studies in mouse models *in vivo* revealed a range of immune responses. Notably, ED2 and ED5 epitope-scaffolds showed a graft-specific and relatively higher level of immune response compared to the other designs. In particular, ED2 which is grafted with two different epitopes showed a robust immune response, suggesting inclusion of multiple epitopes in a single design or inclusion of an epitope-scaffold cocktail for improved efficacy. Surprisingly, these epitope-scaffolds showed limited neutralizing activity in sera. However, when we tested the ED2 binding to serum samples from human COVID-19 patients, ED2 showed graft-specific binding. X-ray crystallographic, SAXS, CD, and ELISA results revealed the stabilized and native epitope-mimicking conformation of the epitope scaffolds. Despite the conformational stabilization, the reason for the limited neutralizing activity of the epitope scaffolds is unclear. However, binding to human patient sera indicates that the plausible reason for limited neutralization response may be species variation. Studies from Correia et al.^22^ previously reported the species-dependent neutralization responses. A limitation of our study is that only linear epitopes were included in the design. Conformational epitopes could be included in further optimization and future designs for improved response.

Here, we conceptually demonstrated the applicability of these epitope-scaffolds in detecting epitope-specific antibodies from serological samples. ED2-based ELISA was used to detect IgG antibodies in serum samples from RT-PCR confirmed COVID-19 patients and pre-pandemic samples. Most COVID-19 patients showed strong reactivity against ED2. All pre-pandemic serum samples remained unreactive. Some samples in the COVID-19 group showed low OD readouts similar to healthy samples, suggesting a lack of or a weak seroconversion^64,68,69^. Next steps may involve testing cross reactivity with patient samples of other coronaviruses. When available, studies may also include the COVID-19 positive samples with known times from symptom onset for better analysis of seroconversion. Nonetheless, our results confirm the usefulness of these epitope-scaffolds for ELISA.

We also show the extension of the conventional detection assays to a user-friendly, magnetic nanoparticle-based ELISA approach suitable for a rapid detection of epitope specific antibodies by coupling epitope-scaffolds to magnetic beads. Examples are shown to highlight the utility of epitope-scaffolds; assay optimization may be required before adopting it to specific needs. The magnetic nanoparticle-based ELISA assays are simple to perform and very robust after optimization. In addition, when the designed epitope-scaffolds are used as immunogens, these assays complement and can be a useful tool to study their effect and to monitor the general immunological status of the population.

In summary, using our established computational approach, we grafted conserved, non-RBD epitopes from SARS-CoV-2 spike protein onto several acceptor scaffold proteins to create epitope-scaffolds. Our results provide important methodology for targeting conserved epitopes of SARS-CoV-2 and an immune-focused approach in pan-sarbeco and universal CoV vaccine design. To our knowledge, this is the first study to design immunogens targeting conserved, non-RBD epitopes of SARS-CoV-2 using a grafting approach. The SARS-CoV-2 immunogens designed in this study show promising results and are worthy of further optimization and development.

## Acknowledgement

The following reagent was deposited by the Centers for Disease Control and Prevention and obtained through BEI Resources, NIAID, NIH: SARS-Related Coronavirus 2, Isolate USA-WA1/2020, NR-52281. We acknowledge support from Huck Institutes for the Life Sciences, National Institutes for Health (1R35 GM134864) and the Passan Foundation. Part of the research reported in here was supported by SIG S10 of the National Institutes of Health under award number S10-OD028589-01 to Dr. Neela Yennawar and by R35GM139587 (to K.A.A). We thank Julia Fecko and Dr. Hemant Yennawar from the X-ray crystallography facility at Pennsylvania State University for obtaining the CD spectroscopy and X-ray crystallography data. We thank Royden Clark for the contribution in performing simulations. Brendan Rackley passed away during this study. We dedicate this paper to his memory.

## Author’s contribution

N.V.D. conceived the idea and directed the project. Y.L.V. initiated the project. Y.L.V. carried out computational design of the proteins, constructs design, proteins expression and binding studies. Y.L.V purified the proteins with the support from B.H. B.R. carried out the DMD simulations with the support from J.W. N.C. performed immunization studies. A.G. and S.K. performed neutralization studies. N.H.Y. performed the crystallography and SAXS studies. M.C. and K.A.A contributed to ED2-MNP assay development. Y.L.V and N.V.D. wrote the manuscript and all co-authors assisted in refining it.

## Data availability

The data that support the findings of this study are available from the corresponding author upon reasonable request

## Competing interests

The authors declare no competing interests

**Supplementary information is available for this paper**

## Notes

### Competing Interest Statement

The authors have declared no competing interest.

