## Supplementary Material for "Adaptation-proof SARS-CoV-2 vaccine design"

### Supplementary Table 1. Sequences and scaffold PDB IDs of epitope-scaffolds.

| SN | Name | Scaffold PDB | Sequence |
|---|---|---|---|
| 1 | ED1 | 1BZ4 | SGQRWELALGRFWDYLRWVQTLSEQVQEELLSSQVTQELRALMDETMKELKAYKSE LEEQLTPVAEETRARLSKELQAAQARLGADMEDVCGRLVQYRGEVQAMLGQSTKEL RNRLASHLGKLQDRLLRDADDLQKRLAVYQAG |
| 2 | ED1 control | | SGQRWELALGRFWDYLRWVQTLSEQVQEELLSSQVTQELRALMDETMKELKAYKSE LEEQLTPVAEETRARLSKELQAAQARLGADMEDVCGRLVQYRGEVQAMLGQSTEEL RVRLASHLRKLRKRLLRDADDLQKRLAVYQAG |
| 3 | ED2 | 4UOS | GDNFEVDLMLNKMIEEIKKMLEKAIKKVKEMLEKMIKEIKKMLENGEDSEKILKKAK EMAEKILKMVIELAEKILKKAKEMAEKILKKVKELGVDNEEVKKMLEKMIKEIKNML ESAIGKVQDMLEKMIKEIKKMLENGEDSEKILKKAKEMAEKILKMVIELAEKILKKAK EMAEKILKKVKELGVG |
| 4 | ED2 control | | GDNEEVKKMLEKMIEEIKKMLEKAIKKVKEMLEKMIKEIKKMLENGEDSEKILKKAK EMAEKILKMVIELAEKILKKAKEMAEKILKKVKELGVDNEEVKKMLEKMIEEIKKML EKAIKKVKEMLEKMIKEIKKMLENGEDSEKILKKAKEMAEKILKMVIELAEKILKKAK EMAEKILKKVKELGVG |
| 5 | ED3 | 2A3J | SQVVLITNINPEVPKEFLQDLLYNLASSQGDILDIVVDLSDDNSGKAYIVFATQESAQAF VEAFQGYPFQGNPLVITFSE |
| 6 | ED3 control | | SQVVLITNINPEVPKEKLQALLYALASSQGDILDIVVDLSDDNSGKAYIVFATQESAQA FVEAFQGYPFQGNPLVITFSE |
| 7 | ED4 | 2A3J | SQVVLITNINPEVPKEKLQALLYALASSQGDILDIVVDLSDDNSGKAYIVFATQESAQA FVEAFQGYKTPPIKDVITFSE |
| 8 | ED4 control | | SQVVLITNINPEVPKEKLQALLYALASSQGDILDIVVDLSDDNSGKAYIVFATQESAQA FVEAFQGYPFQGNPLVITFSE |
| 9 | ED5 | 2Z15 | SGMQLEIQVALNFIISYLYNKLPRRRVNIFGEELERLLKKKYEYKTPPIKDYKGSGFRCI HIGEKVDPVIEQASKESGLDIDDVRGNLPQDLSVWIDPCEVSYQIGEKGPVKVLYVDD N |
| 10 | ED5 control | | SGMQLEIQVALNFIISYLYNKLPRRRVNIFGEELERLLKKKYEGHWYPEKPYKGSGFRC IHIGEKVDPVIEQASKESGLDIDDVRGNLPQDLSVWIDPCEVSYQIGEKGPVKVLYVDD N |
| 11 | ED6 | 2Z15 | SGMQLEIQVALNFIISYKTPPIKDRRRVNIFGEELERLLKKKYEGHWYPEKPYKGSGFR CIHIGEKVDPVIEQASKESGLDIDDVRGNLPQDLSVWIDPCEVSYQIGEKGPVKVLYVD DN |
| 12 | ED6 control | | SGMQLEIQVALNFIISYLYNKLPRRRVNIFGEELERLLKKKYEGHWYPEKPYKGSGFRC IHIGEKVDPVIEQASKESGLDIDDVRGNLPQDLSVWIDPCEVSYQIGEKGPVKVLYVDD N |
| 13 | ED7 | 1HD1 | KMFIGGLSWDGTKKDLKDYFSKFGEVVDCTLKLDPITGRSRGFGFVLFKESESVDKVM DQKYKTPPIKDIDPKRA |
| 14 | ED7 control | | KMFIGGLSWDGTKKDLKDYFSKFGEVVDCTLKLDPITGRSRGFGFVLFKESESVDKVM DQKEHKLNGKVIDPKRA |

*Grafted regions are highlighted in red
*the sequence of MGHHHHHHGSENLYFQG was added to the N-terminal.

**Supplementary Table 2. Optimized protein expression conditions for epitope-scaffolds.**

| SN | protein | MW (kDa) | Growth condition | Expression |
|---|---|---|---|---|
| 1 | ED1 control | 15.8 | 37 ˚C, 4 h | Soluble |
| 2 | ED1 | 15.8 | 37 ˚C, 4 h | Soluble |
| 3 | ED2 control | 20.7 | 18 ˚C, 16 h | Soluble |
| 4 | ED2 | 20.7 | 18 ˚C, 16 h | Soluble |
| 5 | ED3 control | 8.8 | 18 ˚C, 16 h | Soluble |
| 6 | ED3 | 8.8 | 18 ˚C, 16 h | Soluble |
| 7 | ED4 control | 8.3 | 18 ˚C, 16 h | Soluble |
| 8 | ED4 | 8.3 | 37 ˚C, 4 h | Soluble |
| 9 | ED5 control | 13 | 18 ˚C, 16 h | Soluble |
| 10 | ED5 | 13 | 18 ˚C, 16 h | Soluble |
| 11 | ED6 control | 8.8 | - | Insoluble |
| 12 | ED6 | 8.8 | - | Insoluble |
| 13 | ED7 control | 13 | 18 ˚C, 16 h | Soluble |
| 14 | ED7 | 13 | - | Insoluble |

**Supplementary Table 3. Melting temperatures for epitope-scaffolds.**

| Melting temperatures (˚C) | |
|---|---|
| ED1 | 51.85 |
| ED2 | >90 |
| ED3 | >90 |
| ED4 | >90 |
| ED5 | 46.69 |

**Supplementary Table 4. SAXS structural parameters for ED2**.

| | |
|---|---|
| ***Guinier analysis*** | |
| I(0) ($cm^{-1}$) | 10.1 |
| $R_g$ (Å) | 33.5 |
| q min (Å $^{-1}$) | 0.012 |
| q max (Å $^{-1}$) | 0.038 |
| ***P(r) analysis*** | |
| I(0) ($cm^{-1}$) | 0.02 |
| $R_g$ (Å) | 33.7 |
| $D_{max}$ (Å) | 150.4 |
| q range (Å $^{-1}$) | 0.012- 0.262 |
| Porod volume (Å$^3$) | 41988 |

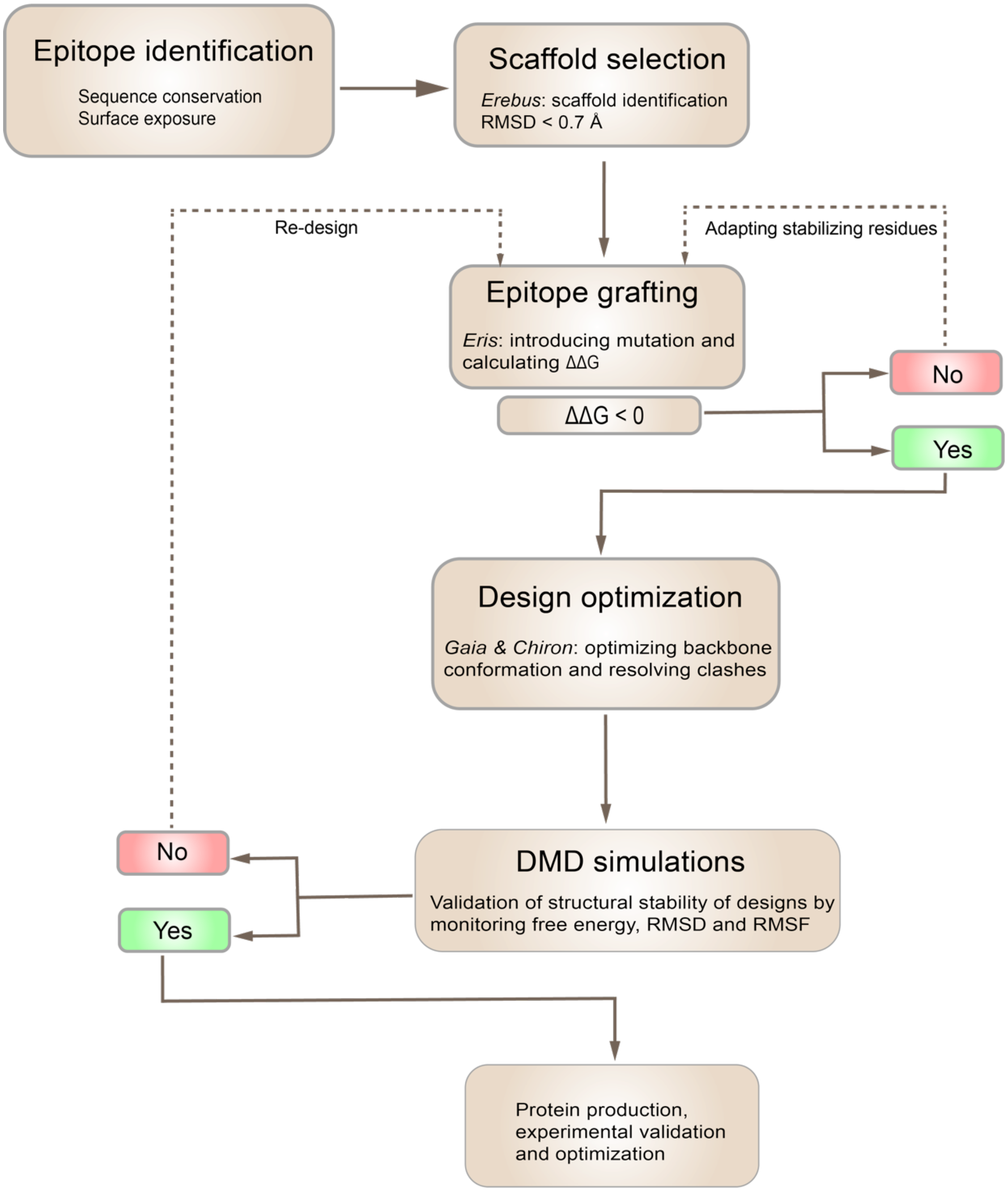

**Supplementary Figure 1.** Flow chart illustrating the key steps in the design of epitope-scaffolds.

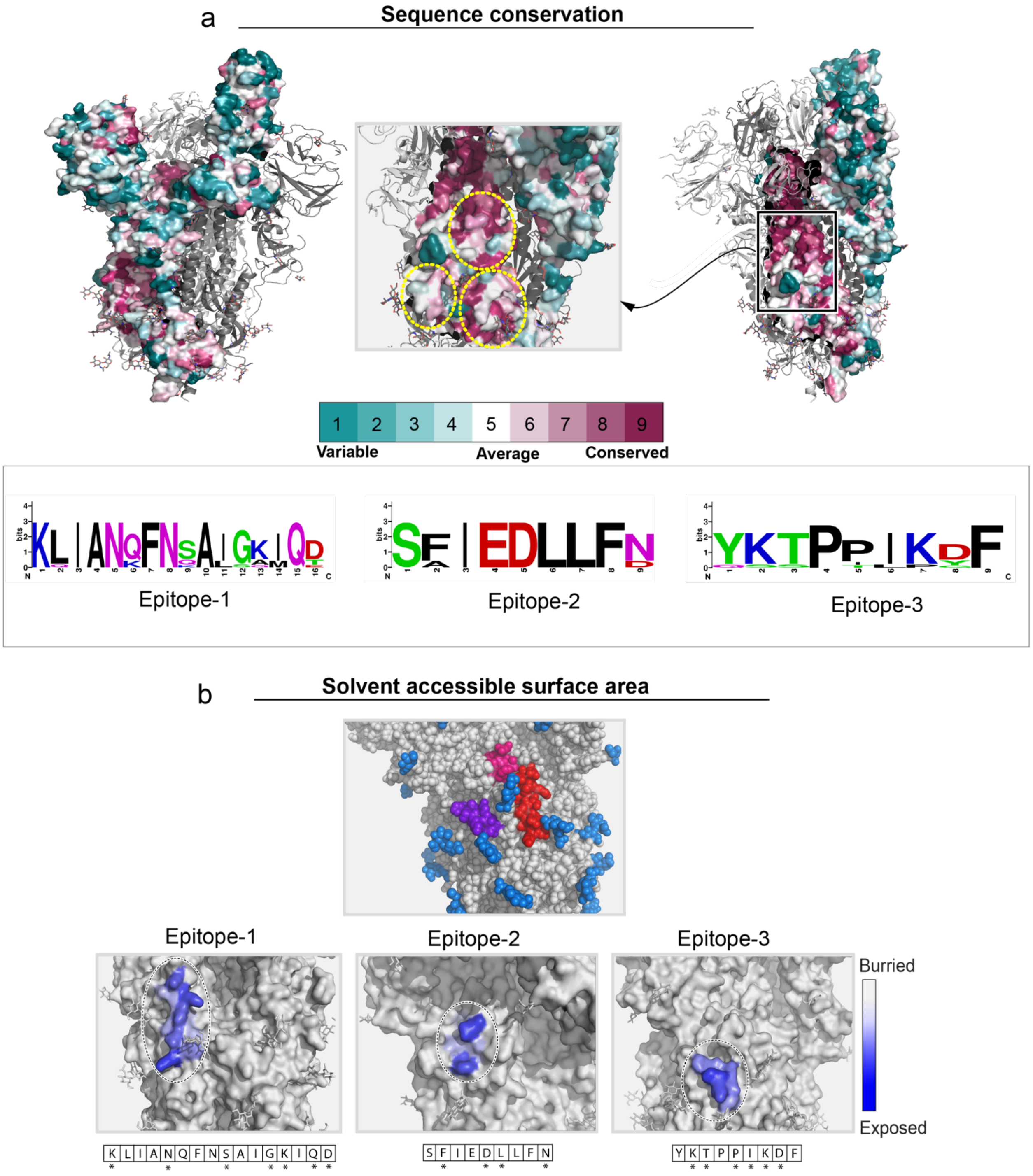

**Supplementary Figure 2. Sequence conservation and solvent accessibility of identified epitope regions**. **a**, Region circled in yellow in the top panel shows the identified epitope regions. Sequence conservation scale is shown below the panel. The lower panel shows the consensus sequence of the epitopes compared to SARS-CoV-2 variants and other coronaviruses. **b,** In the top panel, solvent-exposed residues in three epitopes on spike protein are shown in red, pink, and purple. Bottom panel shows the close-up of solvent-exposed residues in three epitopes. Solvent-exposed residues are also marked with asterisks in the sequence box.

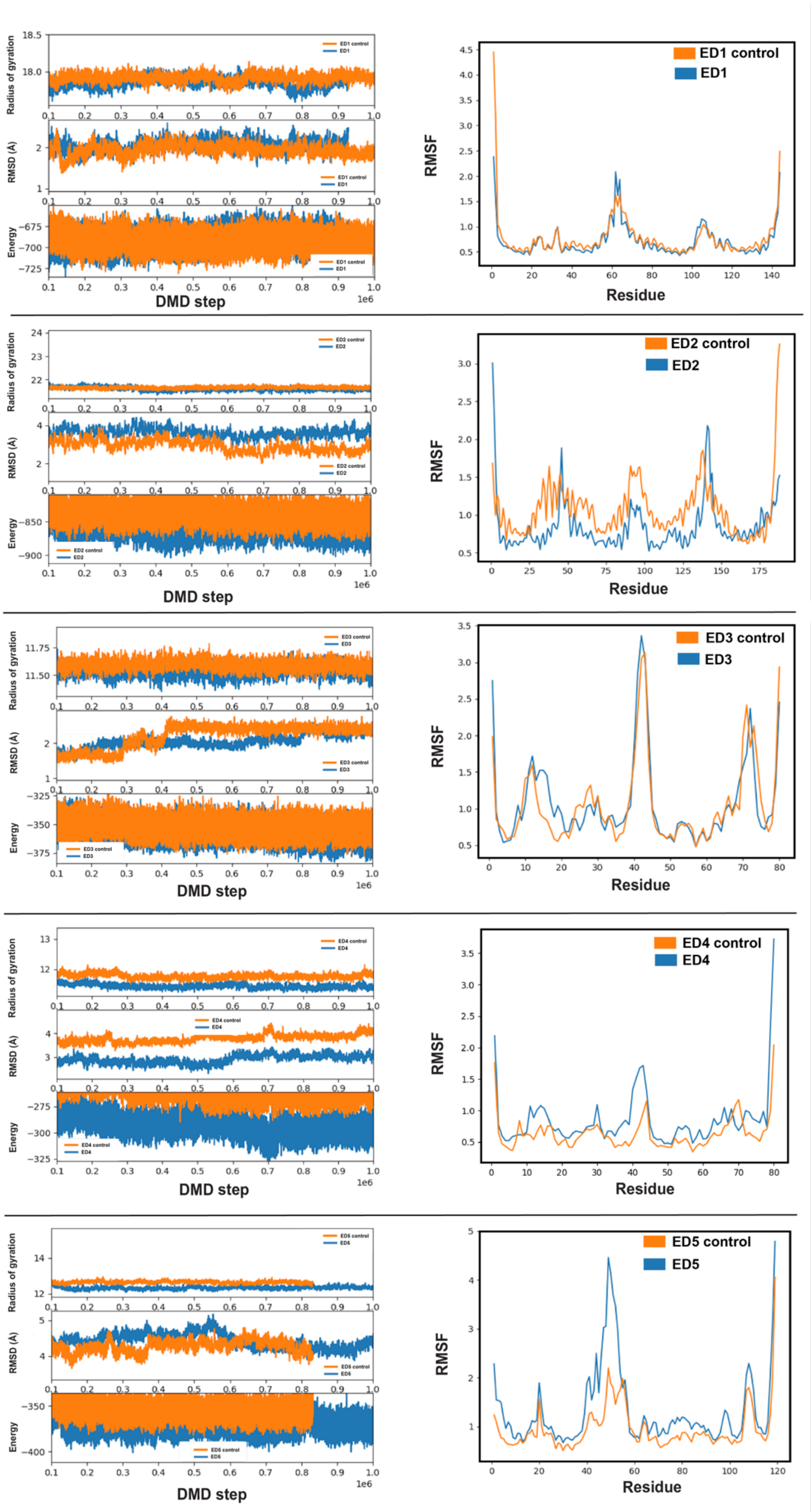

Radius of gyration
RMSD (A)
Energy
ED1 control
ED1
DMD step
1e6
RMSF
Residue
ED2 control
ED2
ED3 control
ED3
ED4 control
ED4
ED5 control
ED5

**Supplementary Figure 3. The conformational plasticity of five best epitope-scaffolds assessed by DMD simulations.** Structural rigidity of grafted epitopes in the epitope-scaffolds was assessed by analyzing RMSD, RMSF, radius of gyration, and energy. For comparison, native scaffold proteins were also included in the simulations. Orange indicates control, non-transplanted scaffolds and blue indicates epitope-scaffolds.

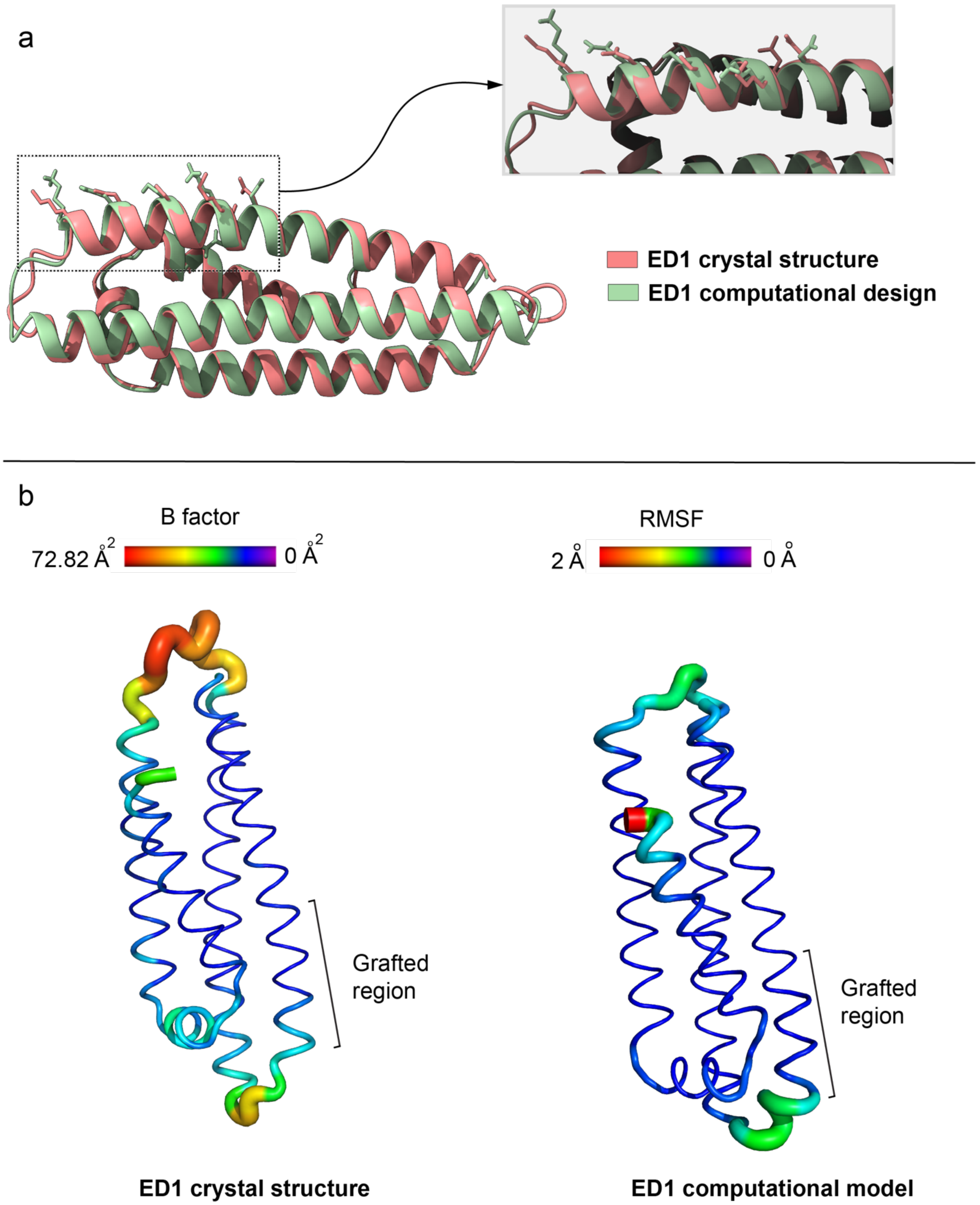

**Supplementary Figure 4. a,** Crystal structure of ED1 aligned to the computational design model, with a close-up view of the grafted region. ED1 is shown in salmon and the computational model is shown in green. Epitope are shown as sticks. **b**, The conformational plasticity of ED1 assessed with B factor (for crystal structure, left) and RMSF (for computational model, right).

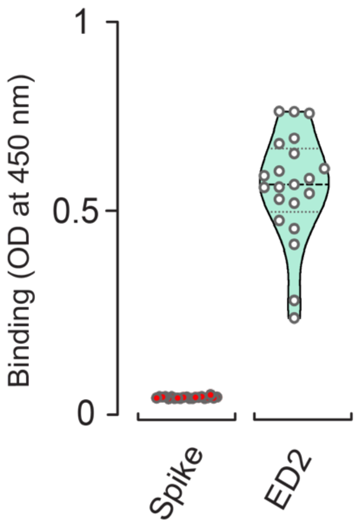

**Supplementary Figure 5.** Comparison of ED2 monoclonal antibodies' (n=21) response to ED2 and spike protein. Data represent violin plots and individual data points.

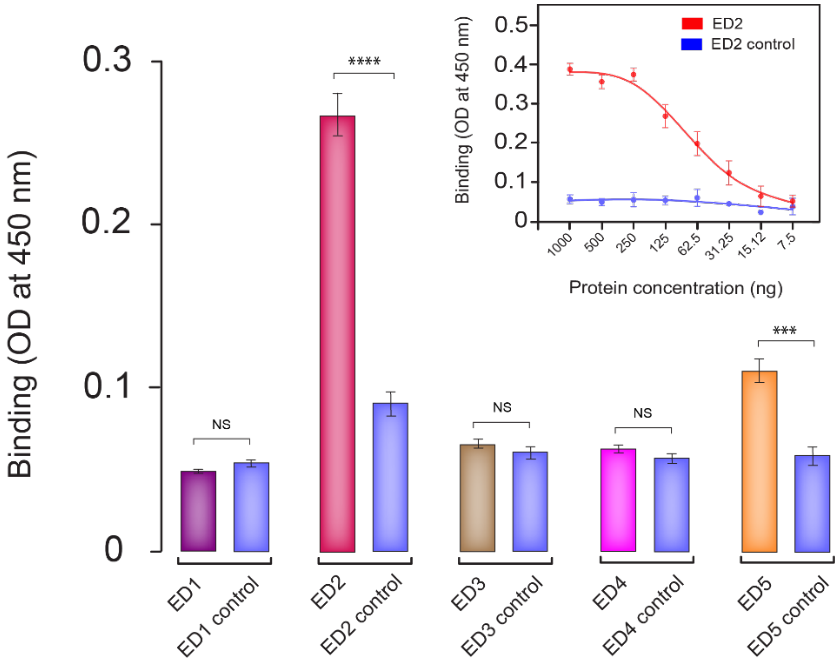

**Supplementary Figure 6. Binding of polyclonal anti-spike antibodies to epitope-scaffolds.** Epitope-scaffolds and controls were tested by ELISA for binding to polyclonal anti-spike antibodies. Inset shows the concentration dependent, graft-specific binding of anti-spike antibody to ED2.

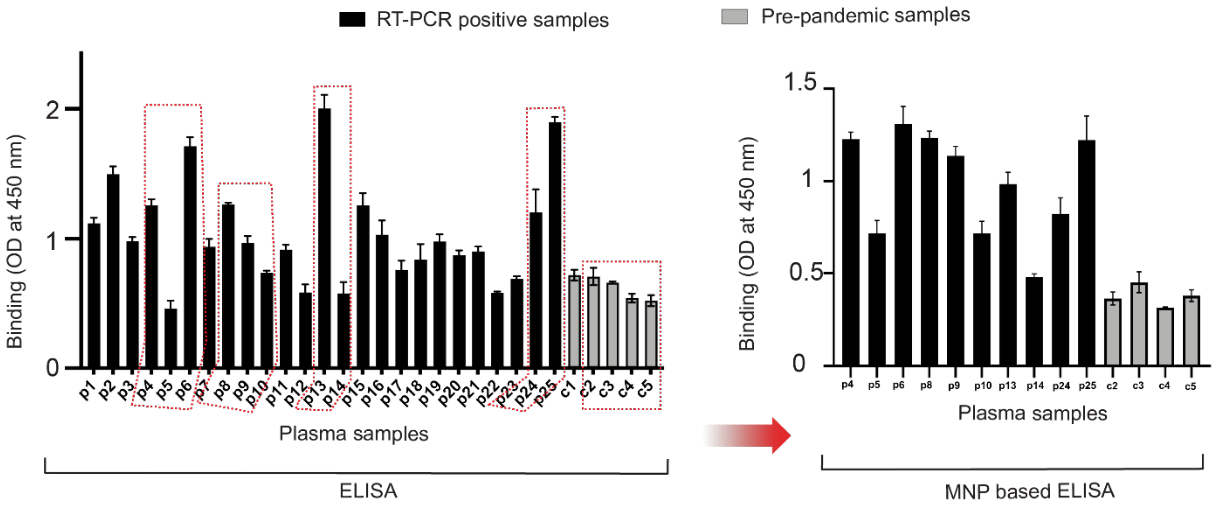

**Supplementary Figure 7. ELISA based on ED2-decorated magnetic nanoparticles.** Anti-epitope-1 IgG antibodies were tested in negative controls and COVID-19 patient samples using ED2 and by employing in-house ELISA and MNP-based ELISA. The red boxes on the left indicate the selected samples for MNP-based ELISA assays.
